# Chitosan oligosaccharide induces plant resistance gene expression in *Pinus massoniana*

**DOI:** 10.1101/2023.01.11.523542

**Authors:** Huayang Yin, Wanlin Guo, Jianmin Fang, Hongjian Liu, Guangping Dong, Xiaojuan Li

## Abstract

Chitosan oligosaccharides are the main degradation products from chitosan or chitin and have been reported to induce resistance to diseases in herbaceous plants like cucumber and *Arabidopsis*. Concomitantly, pine wilt disease is a devastating disease of conifer tree species. Here, we hypothesized that chitosan oligosaccharides induce plant resistance gene (PRG) expression in the woody plant Masson pine, *Pinus massoniana*. Chitosan oligosaccharides were inoculated into *P. massoniana* seedlings and the BGISEQ-500 platform was used to generate transcriptomes from chitosan oligosaccharide-treated *P. massoniana* and control seedlings. A total of 501 differentially expressed genes (DEGs) were identified by comparing the treatment and control groups. A total of 251 (50.1%) DEGs were up-regulated in the treatment relative to the control seedlings and 250 (49.9%) were down-regulated. Inoculation of chitosan oligosaccharide induced the expression of 31 PRGs in *P. massoniana* seedlings and the relative expression levels of six of the PRGs were verified by RT-qPCR. This is the first study to demonstrate that chitosan oligosaccharide induces the expression of PRGs in a tree species. These results provide important insights into the function of chitosan oligosaccharides and further the prospects of developing a chitosan oligosaccharide-based immune inducer for controlling pine wilt disease.

## Introduction

Chitosan oligosaccharide (COS) is a type of polysaccharide, and is the main degradation product from deacetylation and depolymerization of chitosan or chitin via chemical hydrolysis or enzymatic degradation [1]. COS has been shown to induce resistance in *Arabidopsis* to the bacterial pathogen *Pseudomonas syringae* pv. tomato DC3000 and the tobacco mosaic virus [2-3]. Furthermore, fluorinated COS derivatives effectively kill the plant root-knot nematode *Meloidogyne incognita in vitro* and exert excellent control of plant root-knot nematode diseases *in vivo* in cucumbers [4].

*Pinaceae* comprises 11 genera, approximately 230 species, and are widely distributed in the Northern Hemisphere where they play important ecological roles in mountain coniferous forests in northern temperate and subtropical zones [5-6]. The origin of *Pinaceae* can be traced to the Jurassic or even Triassic periods, although the occurrence of modern *Pinaceae* genera primarily occurred in the Early Cretaceous to Tertiary periods [5]. Masson pine (*Pinus massoniana* Lamb.) is a commercially important conifer tree species in South China due to its straight trunk, excellent timber traits, and high oleoresin yields. The species is distributed across 17 administrative provinces from 21°41’N to 33°56’N and 102°10’E to 123°14’E [7].

Pine wilt disease is a devastating disease of conifer tree species and is listed in the phytosanitary directory as an extremely damaging disease of many countries, including in North America, Asia, Europe, and Africa [8-12]. The primary causal agent of pine wilt disease is the migratory endoparasite pine wood nematode *Bursaphelenchus xylophilus* (Steiner et Buhrer) Nickle [13]. *B. xylophilus* can infect about 106 tree species from *Pinaceae* including *P. massoniana, P. thunbergii*, and *P. taiwanensis*. Among the 106 susceptible tree species, 60 can become naturally infected, while 46 can become infected through artificial inoculation [14].

In the present study, we hypothesized that COS helps induce plant resistance gene (PRG) expression in *P. massoniana*, thereby potentially enabling pine trees to develop resistance to pine wilt diseases. COS was consequently inoculated into *P. massoniana* seedlings and the BGISEQ-500 platform was used to generate transcriptomes from COS-treated *P. massoniana* and control samples. Differentially expressed genes (DEGs) between the treated and control groups were specifically analyzed and annotated. Among the DEGs, up-regulated PRGs were especially analyzed. The results from this study provide important new insights into the function of COSs and further the prospects for developing a COS-based immune inducer for controlling pine wilt disease.

## Materials and methods

### Plants and treatments

The field experiments of this study were conducted in July 2021 within the nursery garden of the Anhui Provincial Academy of Forestry. Three-year-old *P. massoniana* seedlings were used in this study. The tops of *P. massoniana* seedling shoots were cut and a 1 mL pipette tip was placed over the notch of the remaining shoot, followed by wrapping the gap between the trunk and the pipette tip with parafilm (Parafilm, Neenah, WI, USA). Then, 1 mL of a 1% COS solution (Aladdin, Shanghai, China) was pipetted into the pipette tip linked to the shoot, while controls were treated with ddH_2_O. Three biological replicates were used for both treatment and control groups and each replicate included one *P. massoniana* seedling. *P. massoniana* seedling needles were collected seven days after treatment and samples were separately flash-frozen in liquid nitrogen, followed by storage at − 80°C.

### Total RNA extraction and cDNA library construction

Total RNA from *P. massoniana* needles was extracted using the CTAB-pBIOZOL reagent and an ethanol precipitation procedure according to the manufacturer instructions. Total extracted RNA was then quality-checked and quantified. cDNA libraries were constructed and sequenced following procedures described by Han et al. and Liu et al. [15-16]. The transcriptome datasets are available in the NCBI database under the accession PRJNA757917.

### Assembly and functional annotation

The SOAPnuke software program (version 1.4.0) was used to obtain clean reads by removing raw reads with adaptors, reads with > 5% ambiguous base calls, and low-quality reads, as previously described [17]. Clean reads were aligned to the reference genome sequence using the Bowtie2 program (version 2.2.5) [18]. The Trinity program (version 2.0.6) was used for *de novo* assembly of clean reads, while the TIGR Gene Indices Clustering Tools (TGICL) was used to cluster transcripts into unigenes and to remove redundant reads. Assembled unigenes were aligned to the National Center for Biotechnology Information (NCBI) non-redundant (nr) protein databases with BLAST (version 2.2.23) [19]. Blast2GO (version 2.5.0) was then used to generate functional annotations of genes [20]. The DIAMOND program (version 0.8.31) was used to compare genes to the Plant Resistance Gene Database (PRGdb) for annotation and identify possible resistance genes by further screening annotation results according to query coverage and amino acid identity values [21-23].

### Differentially expressed gene (DEG) analysis

The software RNA-Seq by Expectation-Maximization (RSEM) (version 1.2.8) was used to calculate the relative transcript abundances in samples as fragments per kilobase of exon model per million mapped fragments values (FPKM) [24]. DEGs were detected with the DEseq2 program based on the negative binomial distribution principle [25-26]. Hierarchical cluster analysis was also conducted with the pheatmap function of the R software program. DEGs were functionally classified based on Gene Ontology (GO) and Kyoto Encyclopedia for Genes and Genomes (KEGG) database annotations. The phyper function of the R software program was used for functional annotation enrichment analysis. Cellular functions with Q-values ≤ 0.05 were generally considered significantly enriched.

### Reverse transcription-quantitative polymerase chain reaction (RT-qPCR)

Six up-regulated plant resistance genes from the DEG analyses were subjected to additional RT-qPCR analysis to validate transcript levels between the oligosaccharide treatment and control groups. RT-qPCR primers were designed based on previously published gene sequences using the Primer3web program (version 4.1.0) (Table 1). The genes were amplified from cDNA templates using the PerfectStart Green qPCR SuperMix (TransGen, Beijing, China) following the manufacturer’s instructions. RT-qPCR was performed on a LineGene 4800 Fluorescent Quantitative PCR Detection System (Bioer, Hangzhou, China). Thermal cycling conditions included 94°C for 30 s, followed by 40 cycles of 94°C for 5 s and 60°C for 30 s. The splicing factor U2af large subunit B-like (U2af)was used as the internal reference gene, and relative mRNA expression levels were calculated using the 2^-ΔΔCT^ method [27]. Student’s paired-sample *t* tests were used to statistically compare transcript levels between treatment and control groups. All statistical analyses were performed using the IBM SPSS Statistics program for Windows (version 26.0; Armonk, NY: IBM Corp.).

**Table 1.**
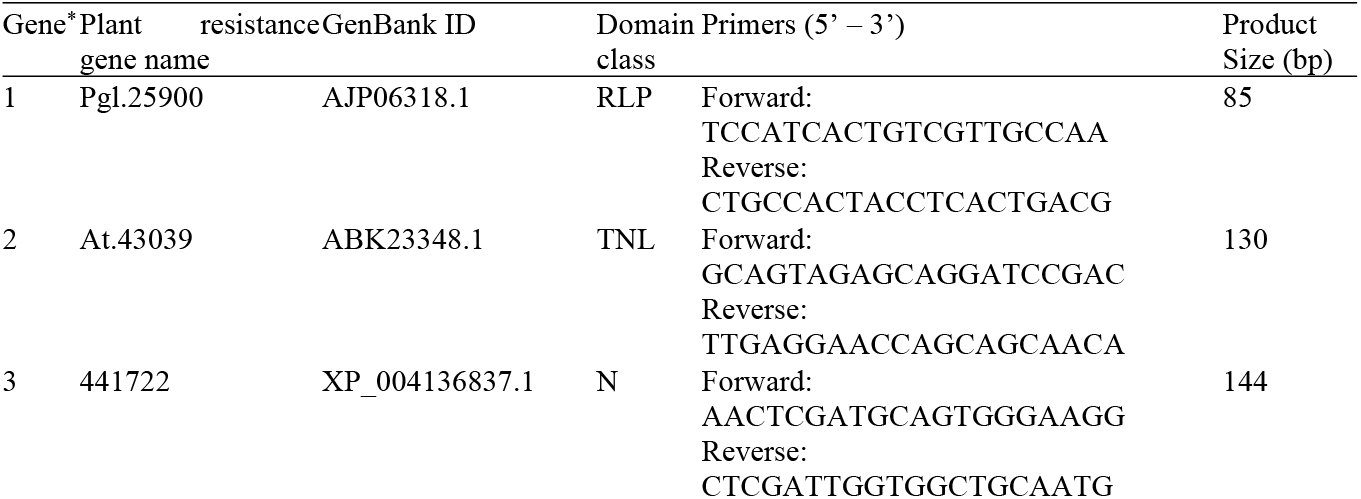
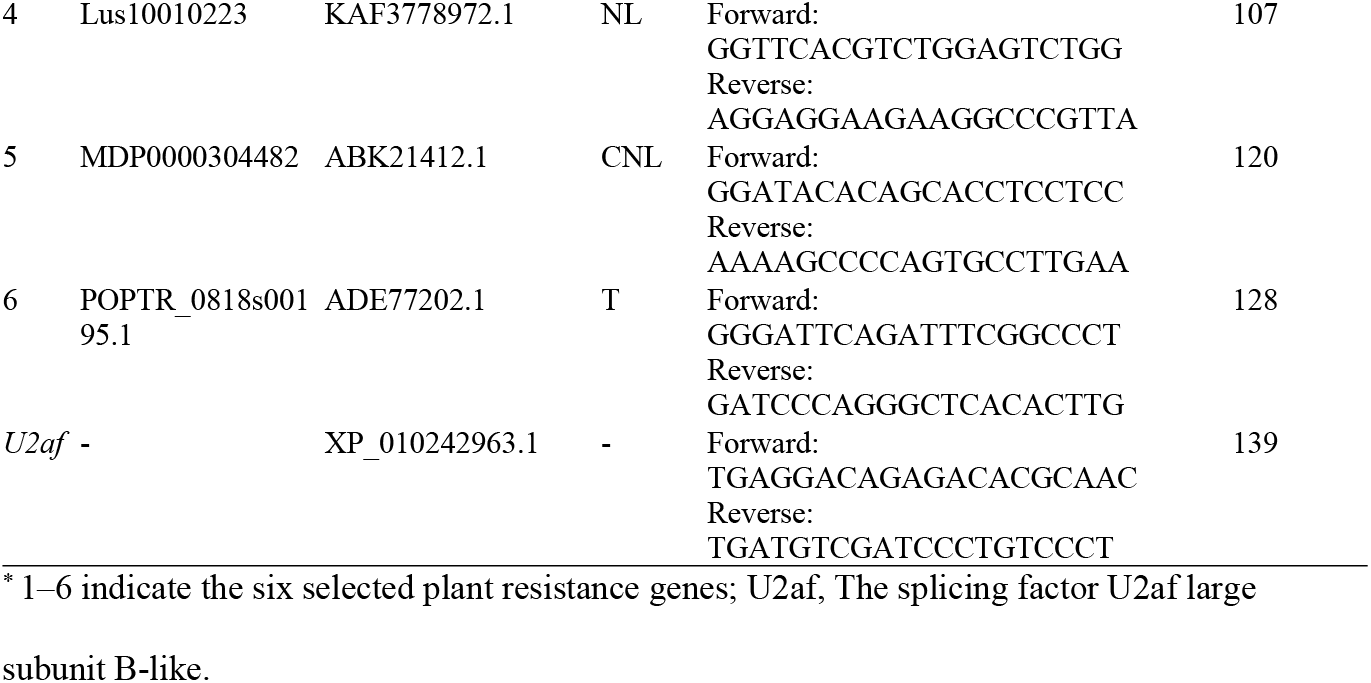
Primers used for RT-qPCR.

## Results

### Transcriptome sequencing and unigene assembly

A total of 273.42 Mbp of raw reads was obtained from transcriptomic sequencing. There were 257.71 Mbp of clean reads after removing adaptors and low-quality reads. A total of 35,493 and 35,272 unigenes were identified via *de novo* assembly from the treatment and control group datasets, respectively. A total of 66,214 all-unigenes were acquired after removing redundant unigenes. The total length of unigenes was 40.11 Mbp for the treatment group, while the mean unigene length was 1,131 bp, the N50 length was 1,791 bp, and the GC percentage was 42.73%. The total length of control group unigenes was 40.79 Mbp, while the mean length was 1,155 bp, the N50 length was 1,794 bp, and the GC percentage was 42.79%.

### Homology analysis and functional gene annotation

Unigenes were aligned against the NCBI nr protein databases for functional annotation using an expected (E)-value threshold of < 10^− 5^ and BLAST comparison. A total of 50,245 (75.88%) of the 66,214 unigenes were annotated in the nr database, while the remaining 15,969 (24.12%) unigenes did not exhibit clear homology with known genes. A total of 22,157 (44.10%) of the unigene sequences were most closely related to proteins from the Sitka spruce (*Picea sitchensis*), followed by *Amborella trichopoda* (3,303, 6.57%), the lotus *Nelumbo nucifera* (1,531, 3.05%), *Cinnamomum micranthum* (1,301, 2.59%) and the loblolly pine, *Pinus taeda* (1,120, 2.23%). The remaining 20,833 (41.31%) transcripts were most closely related to sequences from other plant organisms.

### Functional enrichment of DEGs

Pairwise comparisons enabled the identification of 501 DEGs between the treatment and control groups. In the treatment group, 251 (50.1%) of the DEGs were up-regulated and 250 (49.9%) were down-regulated relative to the control group (Fig 1). The 501 DEGs comprised two major clusters of groups (Fig 2).

**Fig 1.**
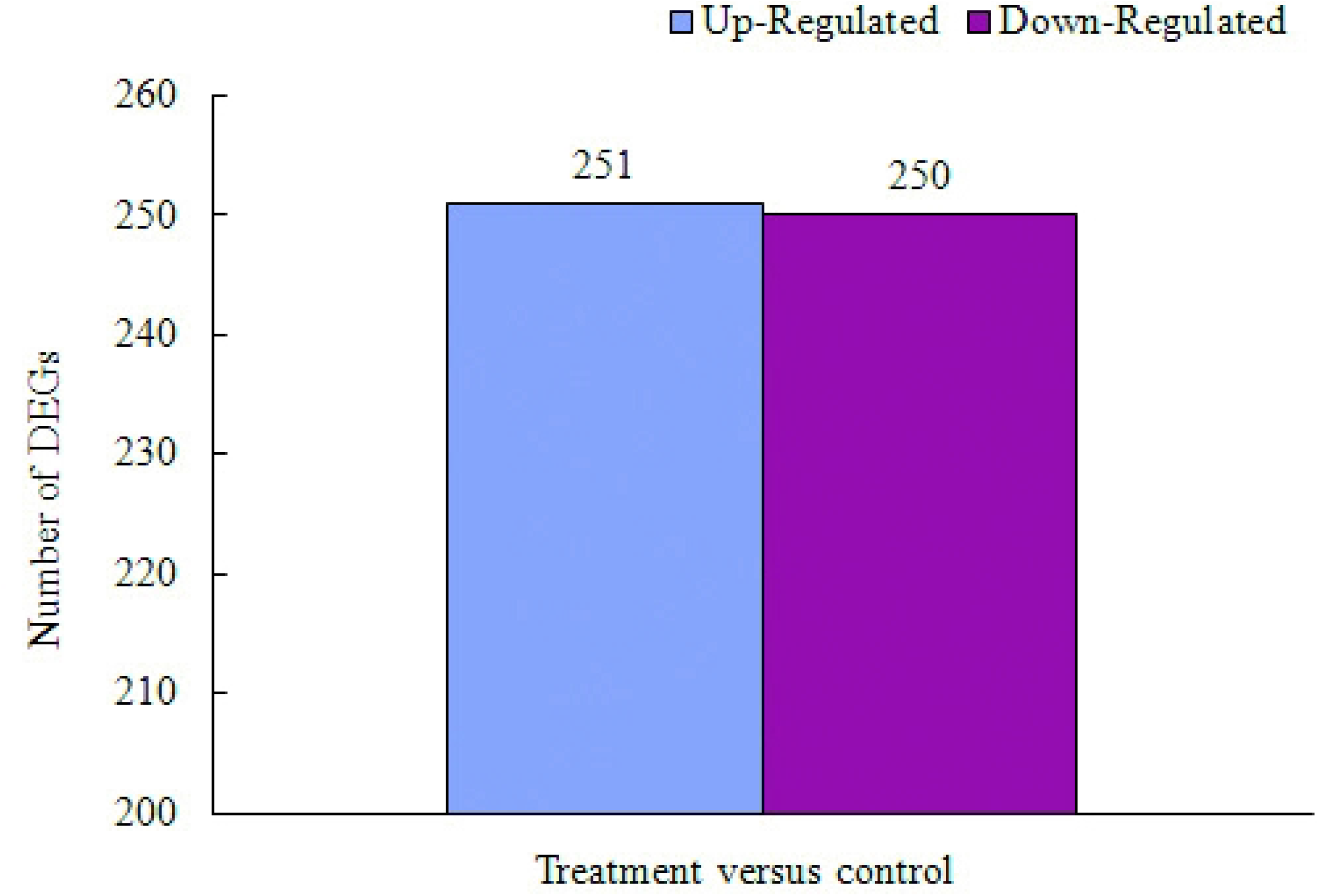
Differentially expressed genes (DEGs) identified by comparison of treatment and control groups. The number above the bar shows the number of up- and down-regulated DEGs between the treatment and control seedling libraries.

**Fig 2.**
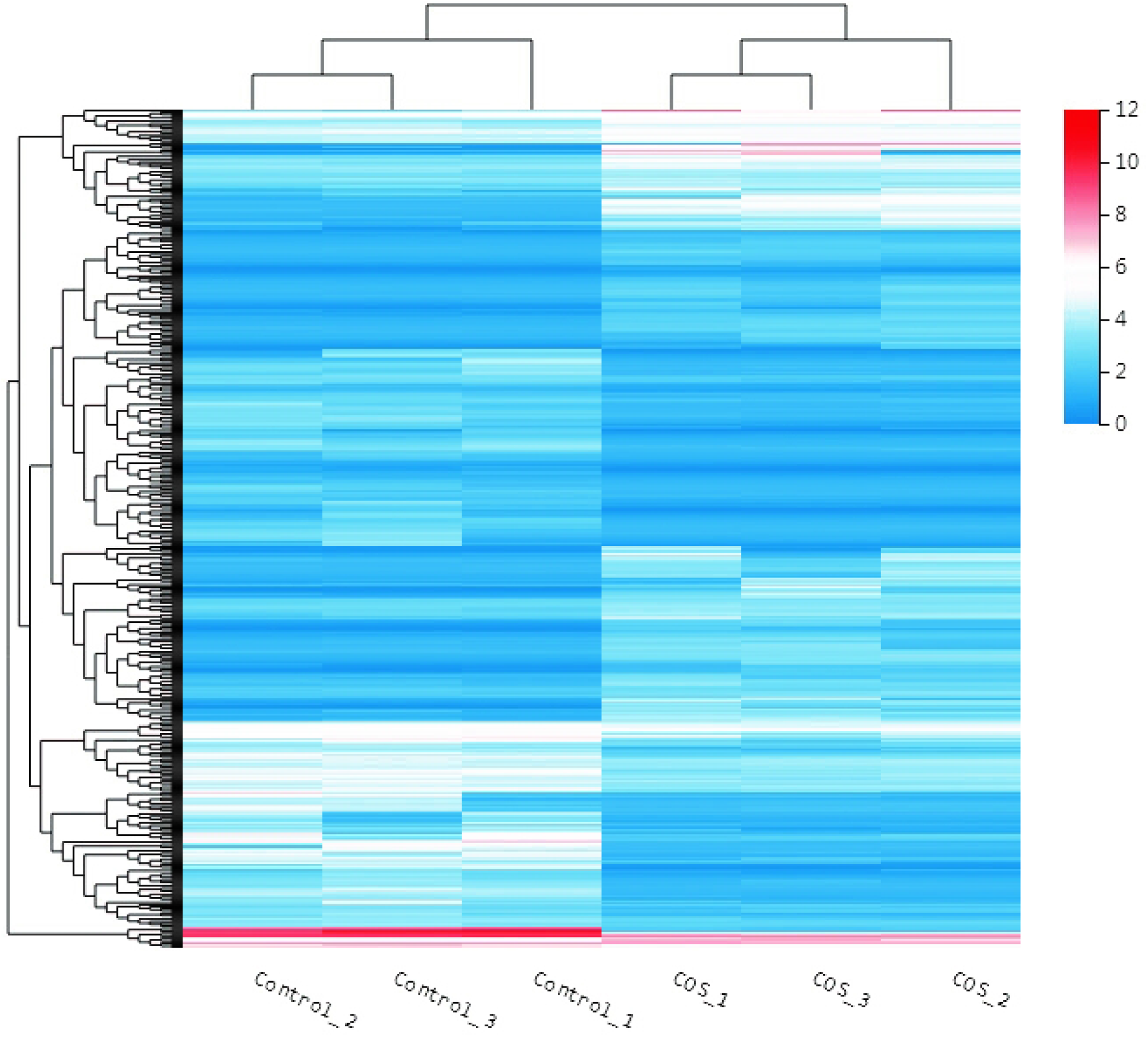
Heatmap of DEGs based on hierarchical cluster analysis. Abundance profiles of all 501 DEGs from the treatment group samples (COS_1, COS_2, COS_3) and control group samples (Control_1, Control_2, Control_3) were identified and compared after normalization using hierarchical cluster analysis. Up-regulated genes are shown in red and down-regulated genes are shown in blue.

GO enrichment analyses indicated that a total of 671 DEGs were significantly enriched. The DEGs were classified into three main GO categories including biological processes, cellular components, and molecular functions that together comprised 31 subcategories. Of the 671 DEGs, 183 (27.27%) were classified within the categories of biological processes, 216 (32.19%) within cellular components, and 272 (40.54%) within molecular functions (Fig. 3). The most highly represented subcategories of the biological processes category were cellular processes (148 genes), metabolic processes (125), and biological regulation (52). The most abundant cellular component terms were cellular anatomical entities (213), intracellular functions (129), and protein-containing complexes (45). Lastly, the most abundant molecular function category annotation groups were binding (162), catalytic activity (162), and transporter activity (24) (Fig 3).

**Fig 3.**
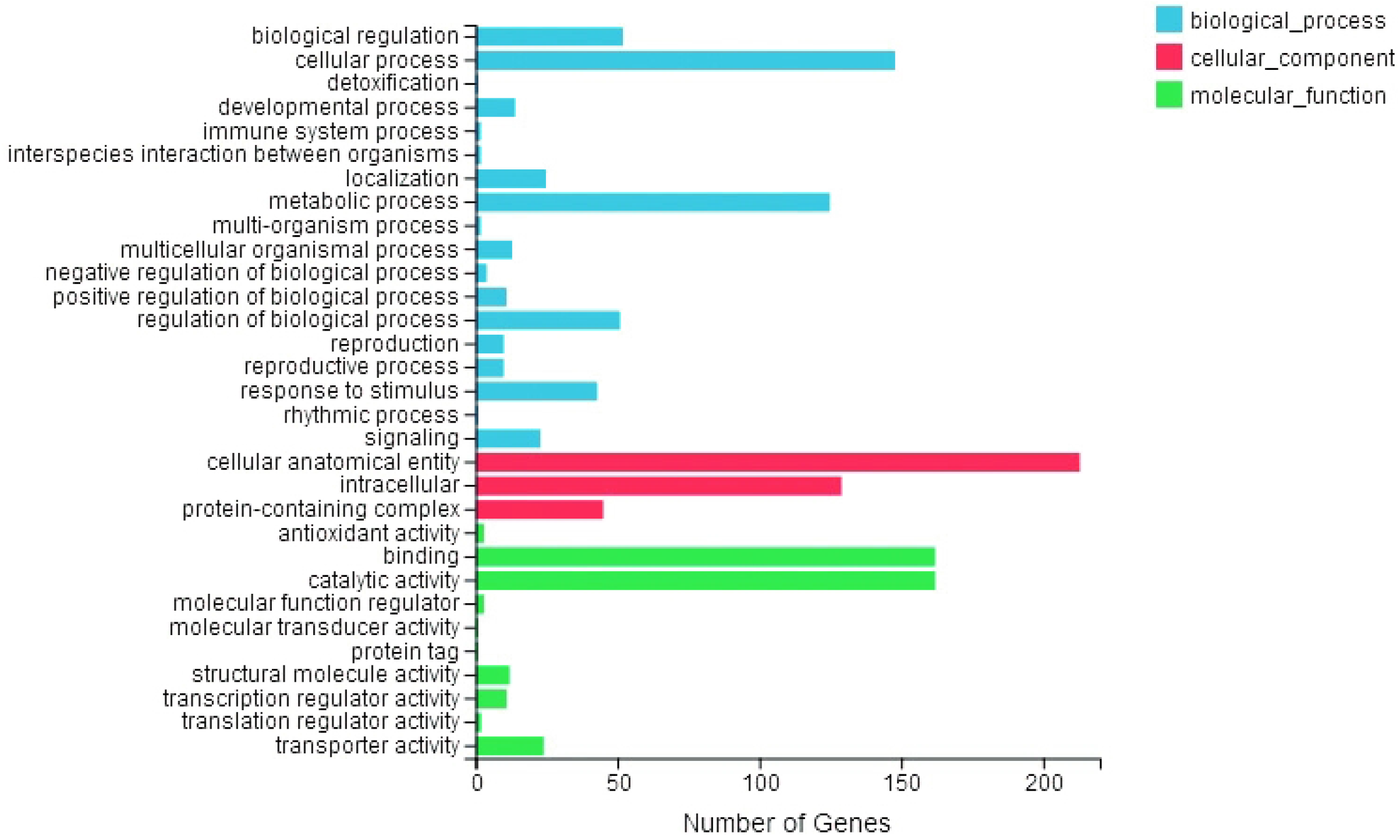
Gene ontology enrichment of DEGs. A total of 671 DEGs were enriched. A total of 235 DEGs were enriched in 19 KEGG pathway subcategories. Of these, 114 DEGs (48.51%) were annotated in metabolism pathways, 63 (26.81%) in genetic information processing pathways, 27 (11.49%) in environmental information processing pathways, 15 (6.38%) in cellular processes pathways, and 16 (6.81%) in organismal system pathways (Fig 4).

**Fig 4.**
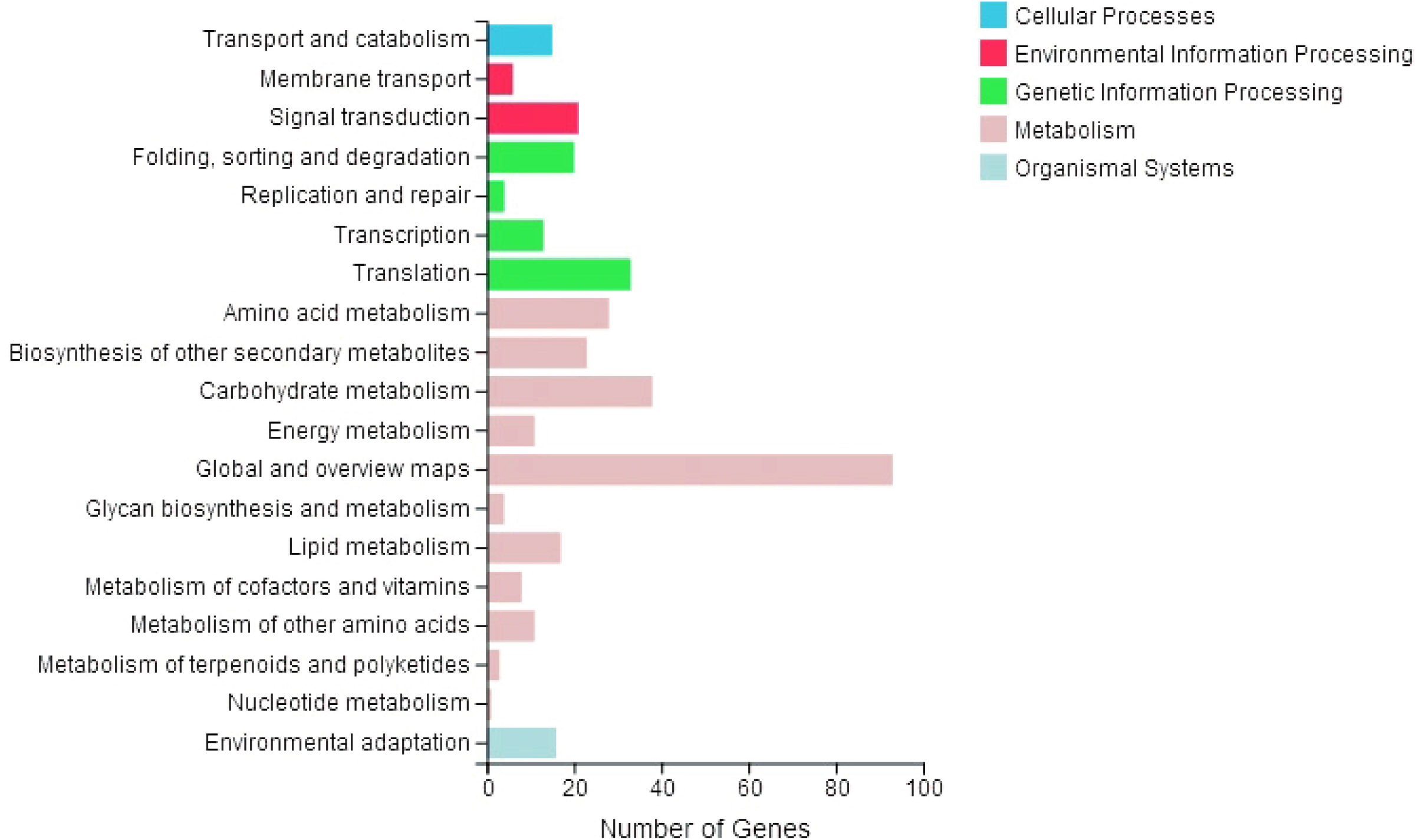
Kyoto Encyclopedia of Genes and Genome enrichment of DEGs. A total of 235 DEGs were enriched.

### Plant resistance genes (PRGs) in *P. massoniana*

Comparison of genes against the Plant Resistance Gene Database yielded the annotation of 6,761 PRGs. Of these, 31 were up-regulated by inoculating COS into *P. massoniana* seedlings (Table 2). Nine up-regulated PRGs belonged to the receptor-like protein (RLP) domain class, six to the Toll/interleukin-1 receptor like protein-nucleotide-binding site-leucine rich re-peat protein (TNL) domain class, and six to the nucleotide-binding site (N) domain class. The nucleotide-binding site-leucine rich re-peat protein (NL) and coiled coil-nucleotide-binding site-leucine rich re-peat protein (CNL) domain classes comprised four and three up-regulated PRGs, respectively. One up-regulated PRG was identified individually for the coiled coil-nucleotide-binding site (CN), the Toll/interleukin-1 receptor like protein (T), and the resistance to powdery mildew locus 8 nucleotide-binding site-leucine rich re-peat protein (RPW8-NL) domain classes. The maximum fold-change of up-regulated PRGs was 8.07 for a PRG with an RLP domain.

**Table 2.**
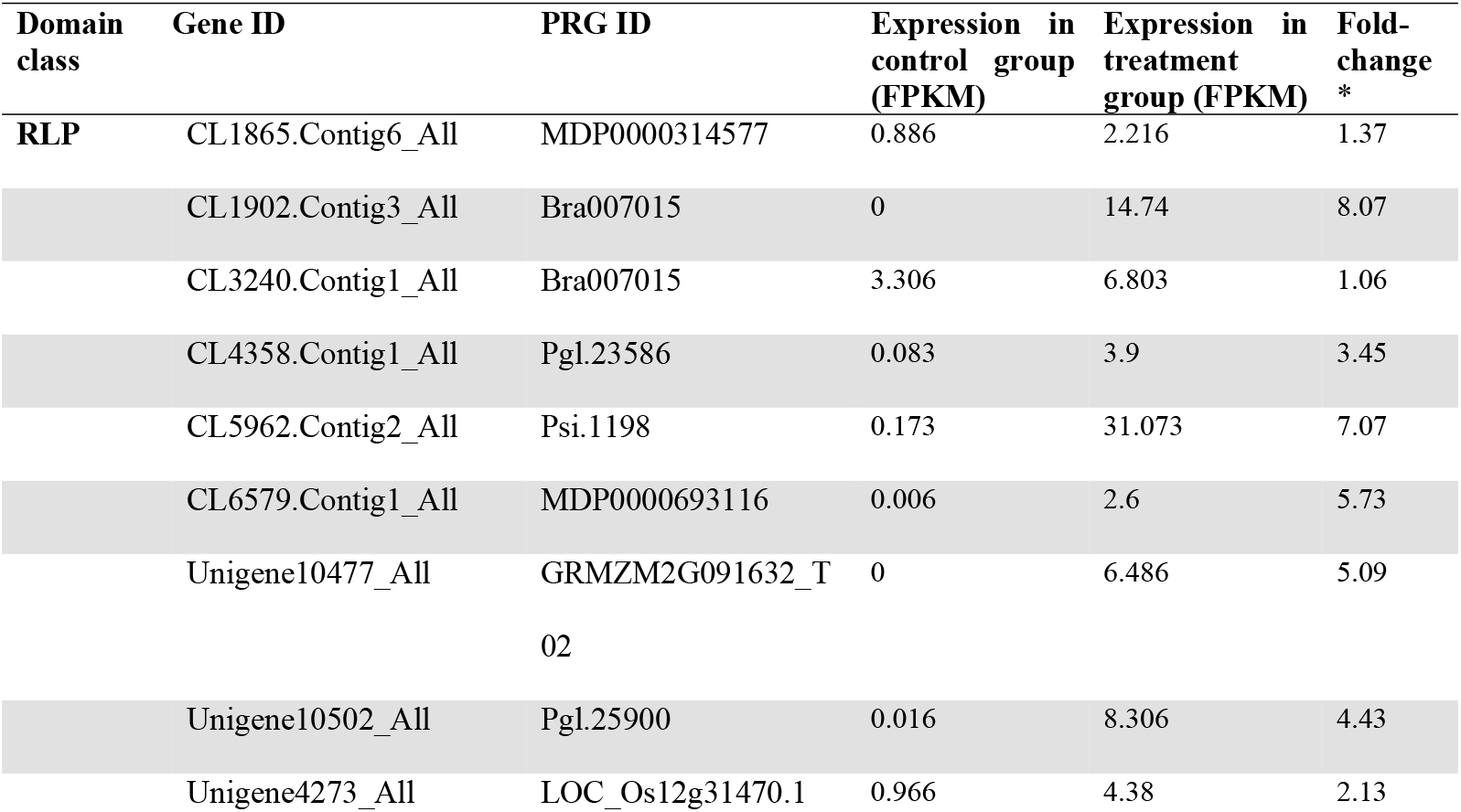
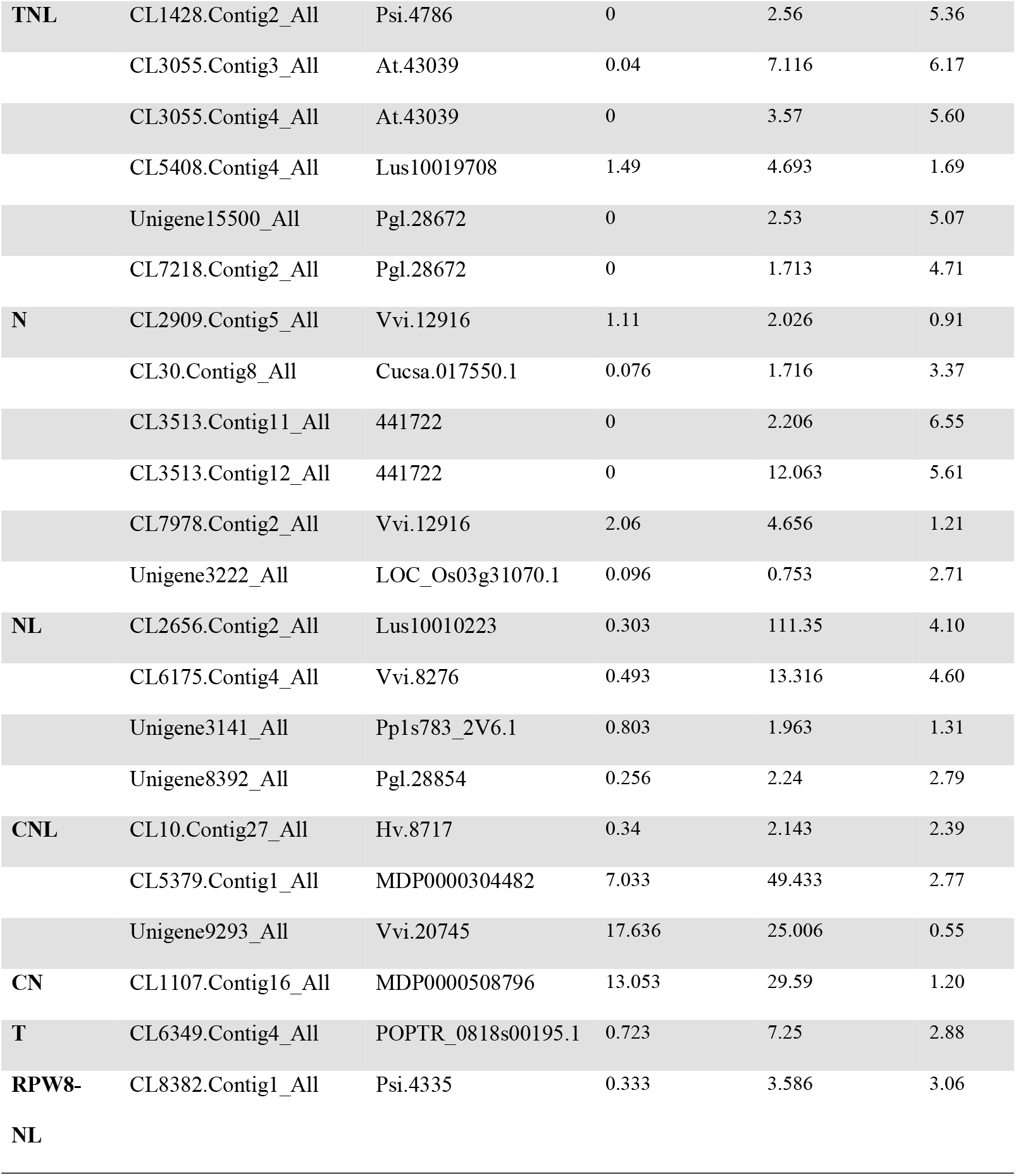
Plant resistance genes (PRGs) that were up-regulated in *P. massoniana* after chitosan oligosaccharide treatment.

### Relative expression patterns of six plant resistance genes after 1% COS treatment

Significant differences were identified in the relative expression levels of four of six up-regulated PRGs between treatments that were selected for additional RT-qPCR validation. Genes significantly up-regulated after 1% COS treatment included Gene1 (*t* = − 6.984; df = 4; *p* = 0.002), Gene3 (*t* = − 5.210; df = 4; *p* = 0.006), Gene4 (*t* = − 4.283; df = 4; *p* = 0.013), and Gene5 (*t* = − 3.357; df = 4; *p* = 0.028; Fig. 5). The remaining two genes were also up-regulated in the treatment group, but not significantly: Gene2 (*t* = − 1.098; df = 4; *p* = 0.334), and Gene6 (*t* = − 1.337; df = 4; *p* = 0.252; Fig 5).

**Fig 5.**
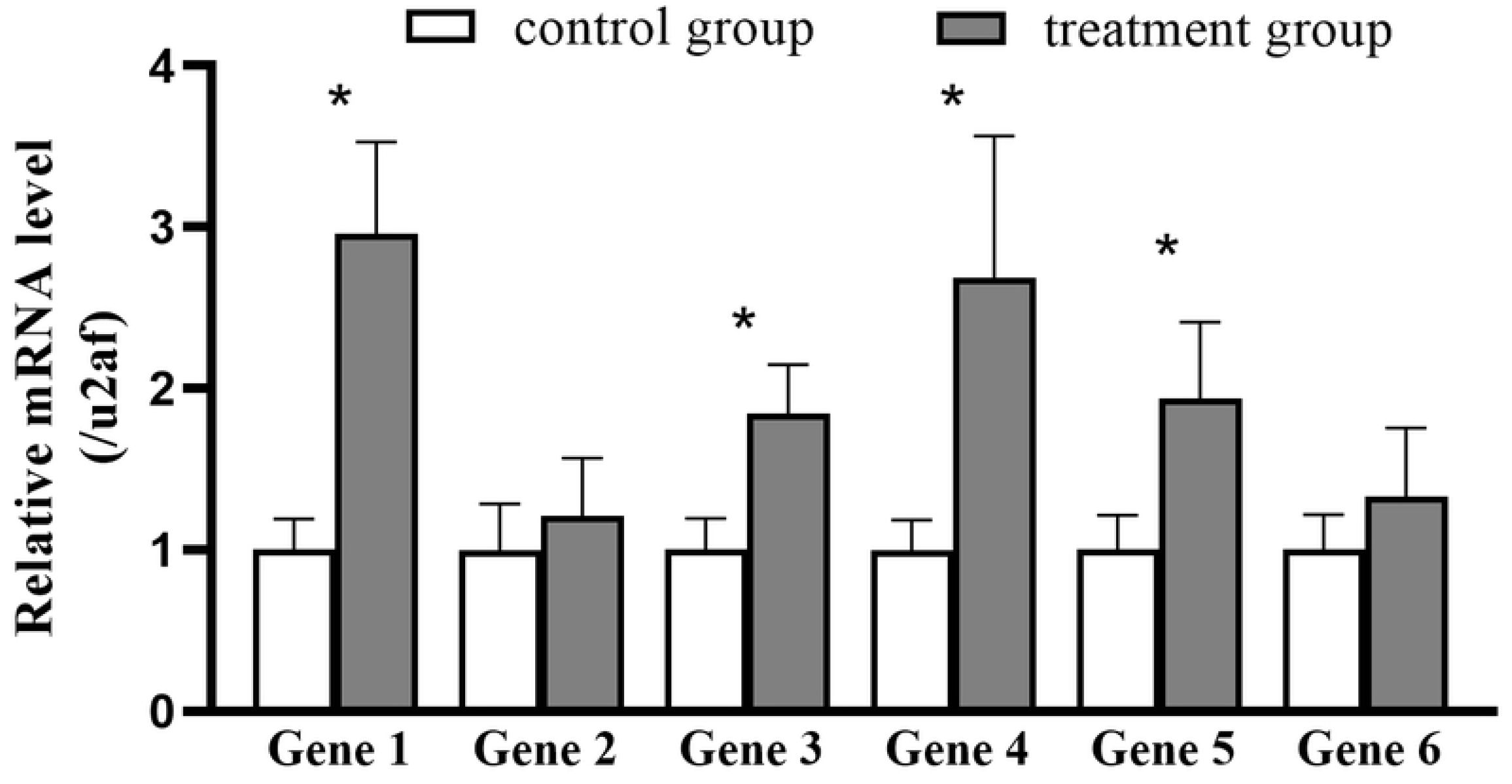
Relative expression levels of six plant resistance genes after 1% chitosan oligosaccharide treatment based on qRT-PCR analysis. The splicing factor U2af large subunit B-like (U2af) was used as the internal reference. Values show the means ± SE of three independent replicates. Asterisks above the bars indicate a significant difference in expression level between the treatment and control groups (*p* < 0.05, pairwise *t* test).

## Discussion

The transcriptome profiles of *P. massoniana* seedlings inoculated with COSs and the control group notably differed. A total of 501 genes were differentially expressed between the treatment and control groups. Compared with the control, 251 (50.1%) of the DEGs were up-regulated and 250 (49.9%) were down-regulated in treated seedlings. GO enrichment analyses revealed that 671 DEGs were significantly enriched. Cellular processes, cellular anatomical entities, and binding were the most abundant subcategories for the biological processes, cellular components, and molecular function categories, respectively, suggesting that cellular process and binding were essential for COS effects on *P. massoniana* seedlings.

A total of 6,761 plant resistance genes from the *P. massoniana* transcriptome were annotated with the Plant Resistance Gene Database. Inoculation of COS in *P. massoniana* seedlings induced the expression of 31 PRGs. The induced PRGs belonged to the RLP, TNL, N, NL, CNL, CN, T, and RPW8-NL domain classes. About one-third of the induced PRGs encoded an RLP domain, indicating that COS-induced PRGs of the RLP domain class likely play important roles in *P. massoniana* pathogen resistance. The relative expression levels of six of the 31 PRGs identified in the transcriptome were validated by RT-qPCR. Among these, four were significantly up-regulated and the remaining two genes were also up-regulated, but not significantly.

PRGs are critical in plant recognition of proteins expressed by specific pathogen avirulence genes that can be functionally classified into five distinct classes according to the presence of specific domains [28-30]. For example, the receptor-like protein (RLP) class includes resistance genes encoding proteins with a receptor serine-threonine kinase-like domain and an extracellular leucine rich repeat (ser/thr-LRR). The TNL class includes proteins with a Toll-interleukin receptor like domain, a nucleotide binding site, and a leucine-rich repeat (TIR-NB-LRR) [21]. The CNL class comprises proteins with a coiled-coil domain, a nucleotide binding site, and a leucine-rich repeat (CC-NB-LRR).

Numerous studies have indicated that PRG expression enhances resistance to diseases in plants. For instance, tomato (*Solanum habrochaites*) *Cf* genes of the RLP class confer resistance to *Cladosporium fulvum* [31]. Further, Whitham et al. described a TNL class resistance gene of tobacco (*Nicotiana benthamiana*) that mediated resistance to the viral pathogen tobacco mosaic virus [32]. Likewise, Sendín et al. reported that transgenic expression of the CNL class gene *Bs2* from pepper (*Capsicum chacoense*) in sweet orange (*Citrus sinensis*) conferred enhanced resistance to citrus canker disease [33]. Nevertheless, further studies are needed to assess whether the PRGs induced by COSs in *P. massoniana* will increase their resistance to pine wilt disease.

## Conclusions

In the present study, transcriptomes from *P. massoniana* seedlings treated with COS and control group were generated and compared. A total of 501 DEGs were identified by pairwise comparisons between the two groups. Among these, 251 were up-regulated in the treatment group in addition to 250 that were down-regulated. Furthermore, inoculation of COS induced the expression of 31 PRGs in *P. massoniana* seedlings. The induced PRGs encoded proteins belonging to the RLP, TNL, N, NL, CNL, CN, T, and RPW8-NL domain classes. In addition, the relative expression levels of six of these PRGs were verified with RT-qPCR. The findings from this study provide novel insights into the development of a chitosan oligosaccharide-based immune inducer for controlling the devastating pine wilt disease and other diseases of tree species.

## Acknowledgments

We thank Shijuan Chen, Xiaosong Yu, and Feng Cheng for their valuable help in conducting field experiments. We thank LetPub (www.letpub.com) for linguistic assistance and pre-submission expert review.

## Notes

### Competing Interest Statement

The authors have declared no competing interest.

